# Dynamic functional connectivity MEG features of Alzheimer’s disease

**DOI:** 10.1101/2023.02.23.529813

**Authors:** Huaqing Jin, Kamalini G Ranasinghe, Pooja Prabhu, Corby Dale, Yijing Gao, Kiwamu Kudo, Keith Vossel, Ashish Raj, Srikantan S Nagarajan, Fei Jiang

## Abstract

Dynamic resting state functional connectivity (RSFC) characterizes time-varying fluctuations of functional brain network activity. While many studies have investigated static functional connectivity, it has been unclear whether features of dynamic functional connectivity are associated with neurodegenerative diseases. Popular sliding-window and clustering methods for extracting dynamic RSFC have various limitations that prevent extracting reliable features to address this question. Here, we use a novel and robust time-varying dynamic network (TVDN) approach to extract the dynamic RSFC features from high resolution magnetoencephalography (MEG) data of participants with Alzheimer’s disease (AD) and matched controls. The TVDN algorithm automatically and adaptively learns the low-dimensional spatiotemporal manifold of dynamic RSFC and detects dynamic state transitions in data. We show that amongst all the functional features we investigated, the dynamic manifold features are the most predictive of AD. These include: the temporal complexity of the brain network, given by the number of state transitions and their dwell times, and the spatial complexity of the brain network, given by the number of eigenmodes. These dynamic features have high sensitivity and specificity in distinguishing AD from healthy subjects. Intriguingly, we found that AD patients generally have higher spatial complexity but lower temporal complexity compared with healthy controls. We also show that graph theoretic metrics of dynamic component of TVDN are significantly different in AD versus controls, while static graph metrics are not statistically different. These results indicate that dynamic RSFC features are impacted in neurodegenerative disease like Alzheimer’s disease, and may be crucial to understanding the pathophysiological trajectory of these diseases.

## 1 Introduction

The human brain can be described as a set of highly dynamic functional networks constructed from a fixed structural network whose fluctuations form the basis for complex cognitive functions and consciousness (Deco and Jirsa, 2012; Shine et al., 2015). Failure of integration within these functional networks may lead to cognitive dysfunction—the cardinal clinical manifestation of Alzheimer’s disease (AD) (Bokde et al., 2009; Knopman et al., 2021; Scheltens et al., 2016). Here, we test the hypothesis that time sensitive descriptions of brain network activity, namely dynamic functional connectivity (FC), are crucial features of functionally relevant alterations in network structure that may underlie AD pathophysiology (Sperling et al., 2010). Although there is a vast literature on static FC and its graph theoretic properties in AD brains, a comparable body of work interrogating the dynamic aspects of FC and its alteration in disease is still lacking. Image data resolution is one obstacle for obtaining convincing evidence that dynamic FC generates strong predictors that distinct between AD and control samples. To date, most dynamic FC studies in AD have focused on low temporal resolution resting state functional magnetic resonance imaging (fMRI) (Schumacher et al., 2019; Sendi et al., 2021; Chumin et al., 2021; Ma et al., 2020; Dautricourt et al., 2022), restricting them only to detect state transitions that may occur in the timescale of seconds. However, micro-states with faster dynamics in the timescale of tens to hundredths of milleseconds are considered the basis for the rapid reorganization and adaptation of the functional networks of the brain (Van de Ville et al., 2010).

Several technical challenges also prevent current studies from demonstrating the utility of dynamic FC features in AD studies. Sliding-window techniques have been commonly applied to extract the dynamic FCs. While the sliding-window method is practically attractive due to its analytical simplicity and easy implementation, it presents several limitations and trade-offs. The temporal resolution of the inferred dynamic FC is inherently limited by the window length and overlap. In practice, this trade-off means that only slow changes in brain dynamics in the time-scale of the window length can be detected or tracked. Furthermore, in almost all current implementations, the sliding-window width is typically pre-specified and is not adaptable to the signal statistics or noise (Jiang et al., 2022), and hence the reliability and reproducibility of dynamic FC patterns are still a challenge (Filippi et al., 2019). Therefore, more comprehensive statistical models are required to extract the dynamic FCs (Filippi et al., 2019). Last but not least, the sliding window approaches typically use K-means clustering on time-resolved FCs to determine the discrete states encompassed by the dynamic FCs. Unfortunately, the performance of K-means clustering suffers from the curse of dimensionality and can be distorted when clustering high-dimensional FCs (Sun et al., 2012).

In the current study, we address these challenges by adopting recent advances in model-based analysis of time-varying FC, and apply them to interrogate the role of dynamic FC in the AD context. We utilize the time-varying dynamic network approach (TVDN) proposed by Jiang et al. (2022) to extract these dynamic FCs from magnetoencephalography (MEG) resting state data in a well characterized cohort of patients with AD and an age-matched control cohort study. MEG has been shown to have good sensitivity to detect early functional changes associated with AD pathophysiology (López-Sanz et al., 2018; Khan and Usman, 2015; Mandal et al., 2018; Maestú et al., 2015). From this high resolution MEG data, TVDN allows us to examine the contributions from temporal and spatial features separately. This is because the TVDN algorithm was designed to ensure that spatial and temporal features from TVDN are not confounded with each other, where the spatial structures arise from the underlying static connectivity, and the temporal parameters describe the dynamic switching between brain networks over time. This is achieved in the TVDN approach by imposing an explicit model of piece-wise constant multivariate signal generation model (see (1) and (2) in Jiang et al. (2022)).

Moreover, TVDN utilizes a data driven dimension reduction and an automated switch detection procedures to capture the dynamic patterns of the FCs. Since this approach requires no clustering of dynamic FCs, it eases the curse of dimensionality and avoids the uncertainties induced by the clustering procedures as those adopted under the sliding window framework. Finally, TVDN selects the model parameters automatically to minimize the uncertainties of the number of switches across independent samples, which generates robust and reproducible dynamic FCs across different datasets.

In Section 4.4 we summarize the TVDN model, its assumptions, and briefly describe how they lead to the desirable properties stated above. In Section 2.1 and Section 2.2, we examine the differences between AD and healthy control groups of the features and graph metrics inferred from the TVDN model. We study the contribution of TVDN features on classifying AD and control subjects in Section 2.3. Finally, we evaluate the sensitivity and specificity of using the TVDN features to predict AD and control classification and compare with benchmark methods in Section 2.4. Using these analyses we show that certain dynamic FC features, including the number of brain state switches, the number of resting state networks, the relative importance of the resting state networks, and a spatial distribution of the resting state networks, are critical for correctly distinguishing AD patients from healthy controls. Our results particularly highlight the importance of dynamic graph metrics over their static counterparts - cementing dynamicity of FC as a key correlate of the disease. We discuss the results and illustrate possible use cases in Section 3. All the technical details are presented in Section 4.

## 2 Results

We implement TVDN on the MEG datasets from 88 AD patients and 88 age-matched healthy control group. All AD patients met the diagnostic criteria for probable AD or mild cognitive impairment due to AD (Albert et al., 2011; McKhann et al., 2011; Jack Jr. et al., 2018). The mean (standard deviation) of the mini-mental state examination score (MMSE) in the AD cohort is 22.14(5.58), and that of the clinical dementia rating (CDR) score is 0.87 (0.49). A schematic of the TVDN is shown in Figure 1, including the set of static and dynamic features extracted from TVDN that will be used in the rest of the paper for the purpose distinguishing AD from healthy control. For each MEG dataset, TVDN automatically detects the brain state switches over given time series, which divides the time series into multiple stationary time segments. We obtain the eigenmodes from TVDN, defined as the magnitude of the top *r* eigenvectors of the implied functional connectivity matrix extending across all the time segments. TVDN assumes the set of eigenmodes remains constant across all time segments, and only their relative contributions change over time. Here *r*, the number of eigenmodes, can vary from one subject to another, and is selected so that corresponding magnitude of the eigenvalues comprises 80% of the total sum of the magnitude of all the eigenvalues. Each eigenmode is a 68 dimensional vector corresponding to 68 cortical regions based on the Desikan-Killiany parcellations (Desikan et al., 2006), and may be thought of as a single resting state network (RSN) that is shared across the time segments. Therefore our RSNs, defined via the TVDN model equation, may or may not correspond to the canonical RSNs one observes via independent component analysis (Yeo et al., 2011). The resulting TVDN scalar features are the *number of eigenmodes* and the *number of brain state switches*. TVDN also provides a spatial feature, the absolute weighted sum of the eigenmodes (WRSN), in each stationary segment, which carries the information of both the shared eigenmodes and the segment specific eigenvalues. The WRSN from each time segment represents the state of the brain during specific time intervals, while the time between two switch points characterizes the dwell time of the brain in each brain state, that is the amount of time the brain spent in a state before moving into a new state. We finally average the WRSN across the segments to obtain the *average weighted resting state network* (AWRSN).

**Figure 1:**
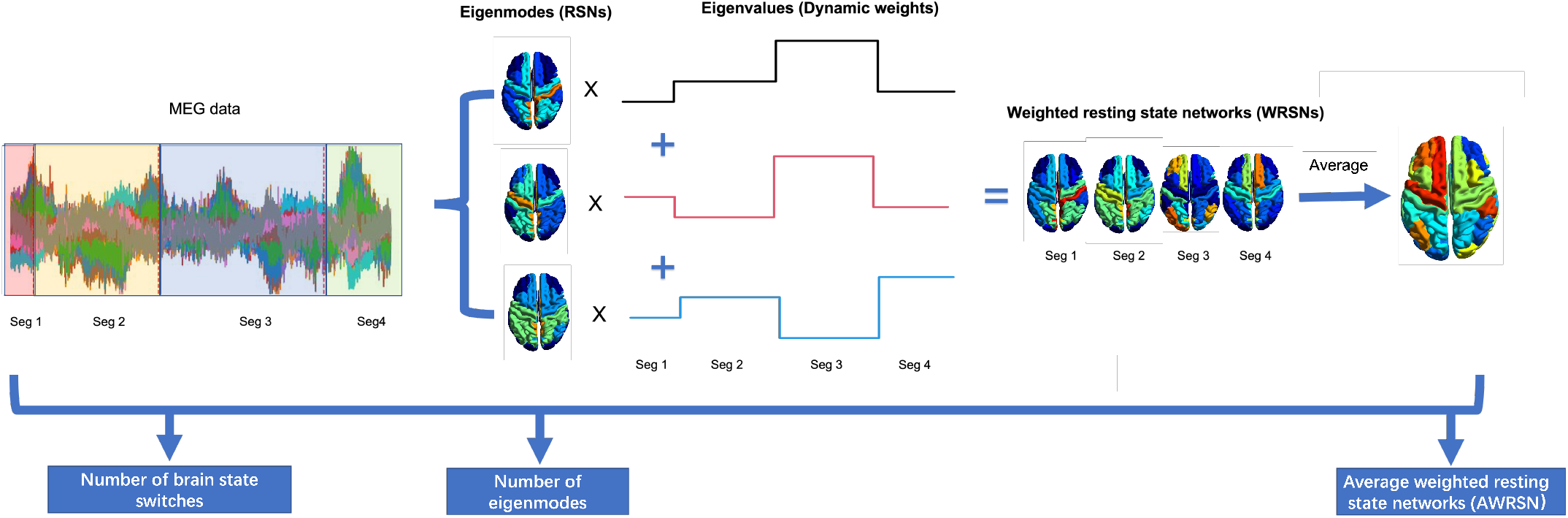
Schematic illustration of TVDN. TVDN extracts spatial features: the number of eigenmodes; temporal features: number of brain state switches; spatial and temporal mixed features: weighted resting state networks (WRSN). The weighted resting state network in each segment is a weighted summation of the eigenmodes with the corresponding eigenvalues as the weights. These are the features used in predictive models described in the rest of the paper.

### 2.1 TVDN scalar features in AD and control groups

We use the above described TVDN method to extract static, dynamic and spatial features from resting state MEG recordings in 88 patients with AD and 88 age-matched-controls participants.

The results in Figure 2 show that the number of eigenmodes are significantly higher in AD group than that in the control group with p-value*<* 0.001 from a student t-test. This is consistent with increase spatial heterogeneity and complexity of dynamic spatial patterns in AD. Despite having an increased number of eigenmodes that commonly used to represent brain states, patients with AD switch less frequently between brain states compared to the healthy controls do on average with p-value=0.007. This is consistent with the observation that the maximal dwell time in a stationary time segment is significantly longer in AD patients than controls with p-value=0.002. These results suggest that AD patients have greater complexity of brain states as represented by higher number of eigenmodes, although they are significantly less active in brain state switches.

**Figure 2:**
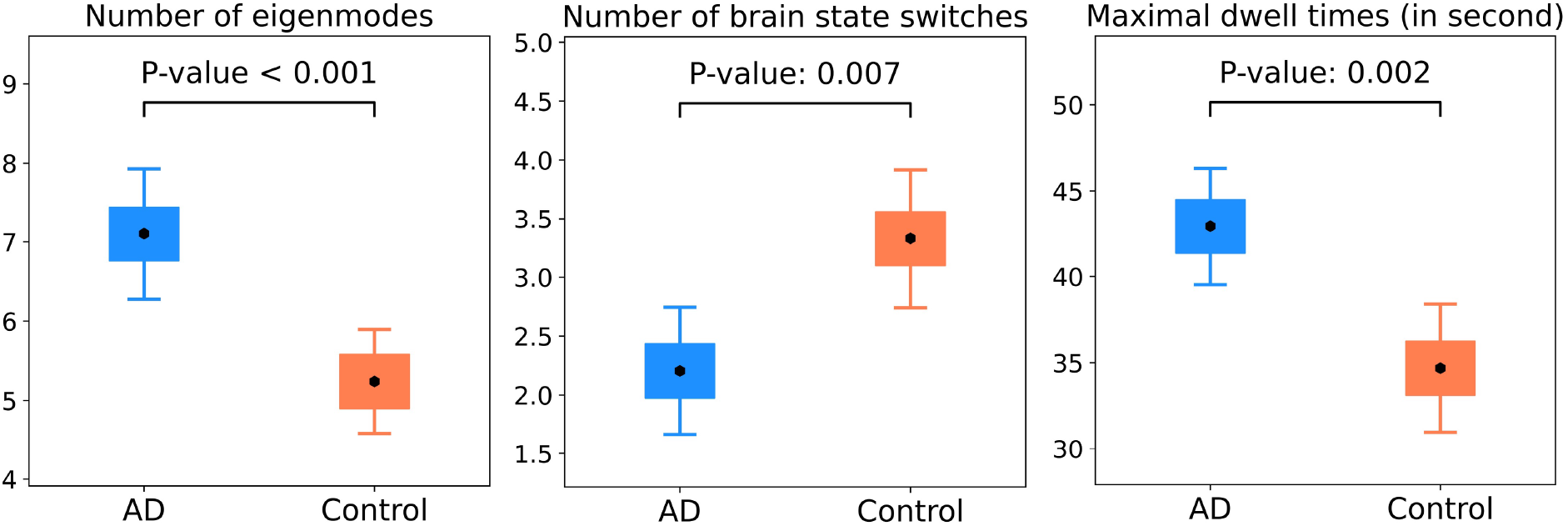
The comparisons between AD and control groups of the static features. Left: number of eigenmodes; Middle: number of brain state switches; Right: maximal dwell times. Mean and 95% confidence intervals are shown and the p-values are from the student t-tests.

### 2.2 Graph metrics of dynamic functional connectivity

We quantify the graph structure of the dynamic FCs using the following graph metrics computed from the brain networks represented by the TVDN connectivity matrices: path length *L*, representing the average shortest length of the path that goes from one region to the other; and modularity *Q*, where a brain network with a higher modularity has denser connections within its subdivisions on average (Stam et al., 2009; Wang et al., 2013; Baniqued et al., 2018). *L* and *Q* are two important graph metrics which measure the *integration* and *segregation* of a brain network (Cohen and D’Esposito, 2016), respectively. Specifically, we compute these graph metrics for all dynamic segments of the data. For each subject, we first obtain the path length and modularity over time denoted by *L*_mean_ and *Q*_mean_, respectively. Furthermore, we extract the graph metrics in the segments with the maximal dwell time denoted by *L*_max_ and *Q*_max_. Moreover, we obtain the variance of the graph metrics over time denoted by *L*var and *Q*var. For a comparison, we summarize the graph metrics of the static FC from the network diffusion model (ND) proposed in Abdelnour et al. (2014), denoted by *L*_static_ and *Q*_static_, where the ND model is a reduced form of the TVDN model when assuming the FC is static. We then compare the properties of these metrics in AD and control cohorts.

Distributions of the variances of the graph metrics over time are shown in Figure 3 (a), which indicates that the path length and modularity from the control cohort have significantly higher variability than those from the AD cohort do after adjusting for the multiple comparison with p-value*<*= 0.025 (0.05*/*2). Therefore, the variations of the graph metrics contain dynamic information that distinguishes AD and control samples. Distributions of the average graph metrics over time are shown in Figure 3 (b), which suggests that the mean of the average modularity is significantly different in the AD and control groups with p-value*<*= 0.047. However, after adjusting for the multiple comparison, none of the average graph metrics is significantly different in the two groups. Distributions of the graph metrics from the segments with the maximal dwell time are shown in Figure 3 (c), which suggests that none of those graph metrics is differentiable in the AD and control groups. Consistent with the results from the segments with the maximal dwell time, the static graph metrics in Figure 3 (d) also do not show between group difference.

**Figure 3:**
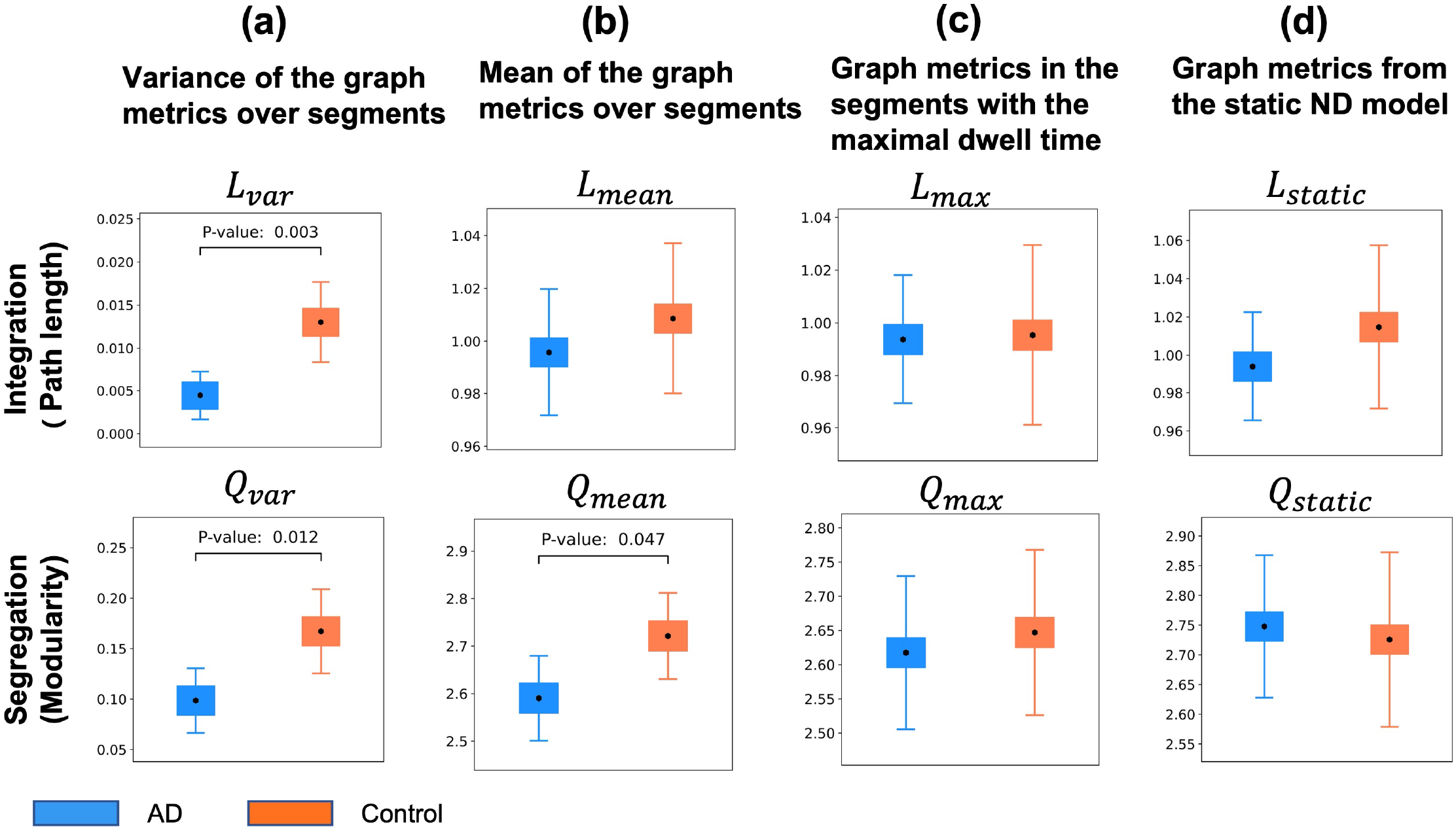
The distributions of the graph metrics extracted from the TVDN dynamic functional connectivity in AD and control groups. (a) The variance of the graph metrics across different segments. (b) The mean graph metrics over the segments. (c) The graph metrics in the segments with the maximal dwell time. (d) The graph metrics from the static FC in the network diffusion model. p-value*<*= 0.05 is used as a significance cutoff. Only the significant p-values are shown in the plots. The error bars represent the 95% confidence interval of the means.

### 2.3 TVDN features are highly associated with AD and control classification

We study the effect of TVDN features and graph metrics to classify AD (group 1) and control (group 0) groups through a logistic regression model. The TVDN features include the number of eigenmodes, the number of brain state switches, and the AWRSN, a 68 dimensional vector, representing the weighted resting state networks over 68 brain regions of interest (ROIs). The graph metrics of interest are the static metrics: *L*mean, *Q*mean, and dynamic metrics: *L*var, *Q*var. Here we do not include *L*_max_, *Q*_max_ because they are highly correlated with *L*mean, *Q*mean, and none of them is associated with the AD and control classification as shown in Figure 3. Since some predictors are highly correlated and the number of predictors is large, we add a ridge regularization to the model to ease the collinearity among the predictors. Moreover, we utilize the bootstrap method (Efron, 1979) to construct the 95% confidence interval (CI) and the p-values of the effects from the TVDN features. We use p-values=0.05 as the cutoff to determine significant difference between AD and control groups (Table 1).

**Table 1:** TVDN results from a ridge regularized logistic regression: the effects from the predictors of interest on the AD and control classifications (outcome). The predictors include the number of eigenmodes, number of brain state switches, the dynamic and static graph metrics and the AWRSN over 68 regions (the labels of the ROIs are presented in the table) of interest are presented along with the corresponding 95% CI and p-value. All predictors are all standardized by their sample means and standard deviations. Only the significant spatial features are shown. p-value*<*= 0.05 is used as a significance cutoff.

| TVDN scalar features |  |  |  |
| --- | --- | --- | --- |
| Features | Effect | 95% CI | P-value |
| Number of eigenmodes | 1.093 | ( 0.572, 1.613) | < 0.001 |
| Number of brain state switches | -0.369 | (-0.731, -0.007) | 0.046 |
| TVDN graph metrics |  |  |  |
| Metrics | Effect | 95% CI | P-value |
| $L_{\text{var}}$ | -0.161 | (-0.645, 0.323) | 0.513 |
| $Q_{\text{var}}$ | -0.153 | (-0.478, 0.173) | 0.358 |
| $L_{\text{mean}}$ | 0.196 | (-0.234, 0.626) | 0.372 |
| $Q_{\text{mean}}$ | -0.273 | (-0.620, 0.073) | 0.122 |
| TVDN spatial features (p-value≤ 0.05) | | | |
| Features | Absolute effect | 95% CI | P-value |
| Left rostral anterior cingulate | 0.665 | (0.319, 1.010) | < 0.001 |
| Left fusiform | 0.619 | (0.293, 0.944) | < 0.001 |
| Left lingual | 0.492 | (0.188, 0.797) | 0.002 |
| Left inferior parietal | 0.611 | (0.228, 0.993) | 0.002 |
| Right inferior temporal | 0.463 | (0.153, 0.774) | 0.003 |
| Left parahippocampal | 0.383 | (0.122, 0.645) | 0.004 |
| Left temporal pole | 0.370 | (0.077, 0.664) | 0.013 |
| Right pericalcarine | 0.432 | (0.068, 0.796) | 0.020 |
| Left superior temporal | 0.332 | (0.052, 0.612) | 0.020 |
| Left lateral occipital | 0.446 | (0.065, 0.827) | 0.022 |
| Left inferior temporal | 0.392 | (0.049, 0.734) | 0.025 |
| Left superior parietal | 0.414 | (0.041, 0.786) | 0.029 |
| Right cuneus | 0.361 | (0.016, 0.707) | 0.040 |
| Right inferior parietal | 0.380 | (0.006, 0.754) | 0.046 |

Consistent with the previous group comparison, the logistic regression showed that AD patients have a greater number of eigenmodes (Table 1, positive estimators), and lesser number of brain state switches (Table 1, negative estimator), compared to controls. Next, we examine the regional patterns of the estimated absolute effects from AWRSN (Figure 4 (a)), which shows the AWRSNs at twelve ROIs are significantly different in AD and control groups when adjusting for other predictors in the model. It is worth mentioning that the six graph metrics do not significantly affect the AD and control classification after adjusting for the other predictors.

**Figure 4:**
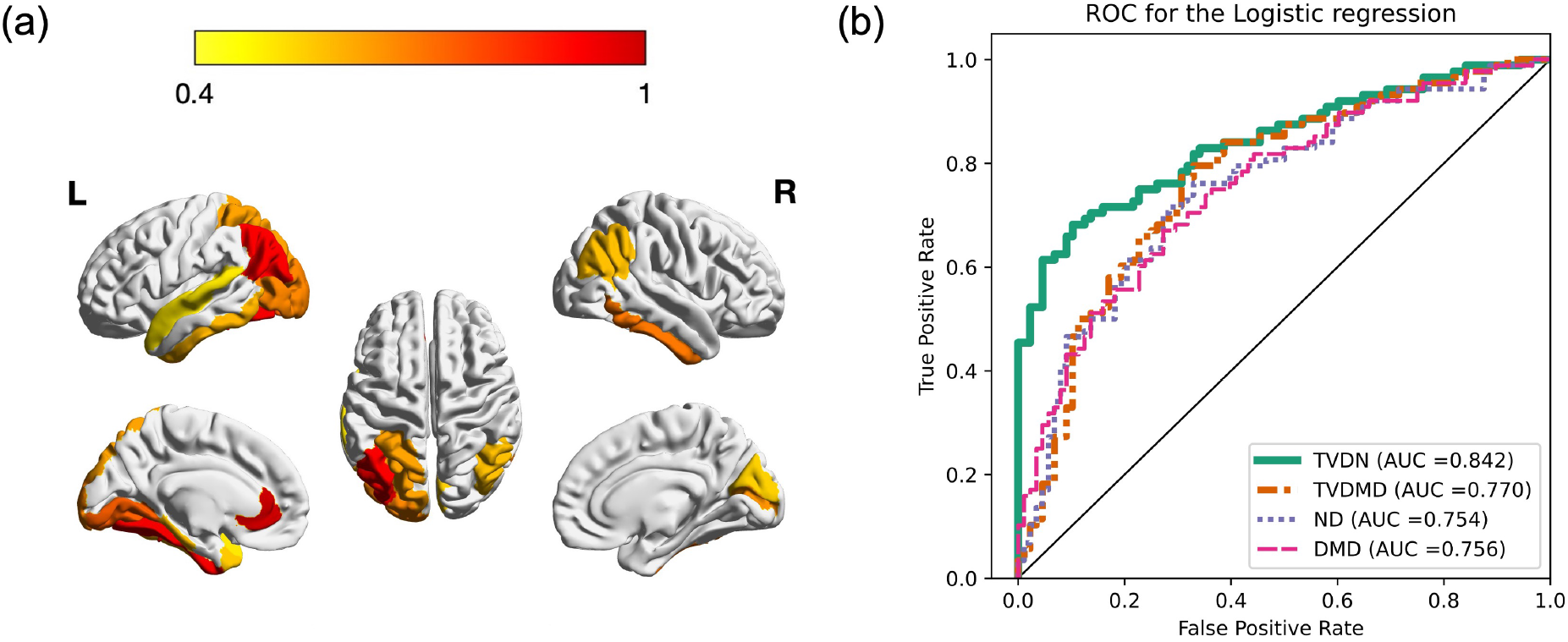
The results from regularized logistic regressions: (a) The estimated absolute effects from AWRSN projected to [0, 1] interval. Only the regions with p-values less than 0.05 are shown in the figure. (b) The ROCs of the leave-one-out prediction results from different models.

### 2.4 Prediction and benchmark comparisons

We perform a leave-one-out (LOO) procedure to examine the accuracy of using significant TVDN features identified in Table 1 to classify AD and control samples. We first use 175 samples to train a ridge regression model and predict the classification of the remaining one sample. We iterate the procedure to predict the classification for all samples and to depict the receiver operating characteristic (ROC) curve in Figure 4 (b). The area under the ROC curve (AUC) of using the features from TVDN is close to 84%, suggesting that the significant TVDN features have good sensitivity and specificity to distinguish AD with control subjects, and hence can be used to improve application considering other more conventional features in future applications (Metz, 1978; Obuchowski, 2003).

We compare the prediction performance of using features from TVDN model to the prediction performances of using the features from two static FC models: dynamic mode decomposition model (**DMD**) (Brunton et al., 2016) and network diffusion model (**ND**)(Abdelnour et al., 2014) and from one dynamic FC model: time-varying dynamic mode decomposition model (**TVDMD**) (Kunert-Graf et al., 2019). DMD assumes that the observed signal follows a multivariate autoregression model, ND links two signals at consecutive times through a differential equation model, while TVDMD is a sliding window based extension of DMD. We extract the predictors including the number of eigenmodes, AWRSN and static graph metrics from the static FC models. We also extract the predictors including the number of eigenmodes, the number of brain state switches, AWRSN, static and dynamic graph metrics from the dynamic FC models. The detailed derivations of the predictors are presented in Appendix 4.5 and 4.6. Similar to those used in TVDN evaluation, we first perform a ridge regression to select important predictors based on their confidence intervals and then utilize an independent ridge regression model on the selected predictors to classify AD and control samples. We depict the prediction algorithms in Section 4.7. In addition to the LOO we performed a five-fold cross-prediction, where we used 80% of samples to train the model and predicted the classifications of the rest 20% samples. All the tuning parameters were tuned based on the training data to ensure no information leaking in the prediction procedure. We show the average AUC and their standard deviations from 10000 five-fold cross prediction experiments in Table 2. The corresponding ROC curves under different models are depicted in Figure 4 (b). The results show that TVDN performs the best among all the methods in classifying AD and control samples. The dynamic FC models perform generally better than the static models in the predictions.

**Table 2:**
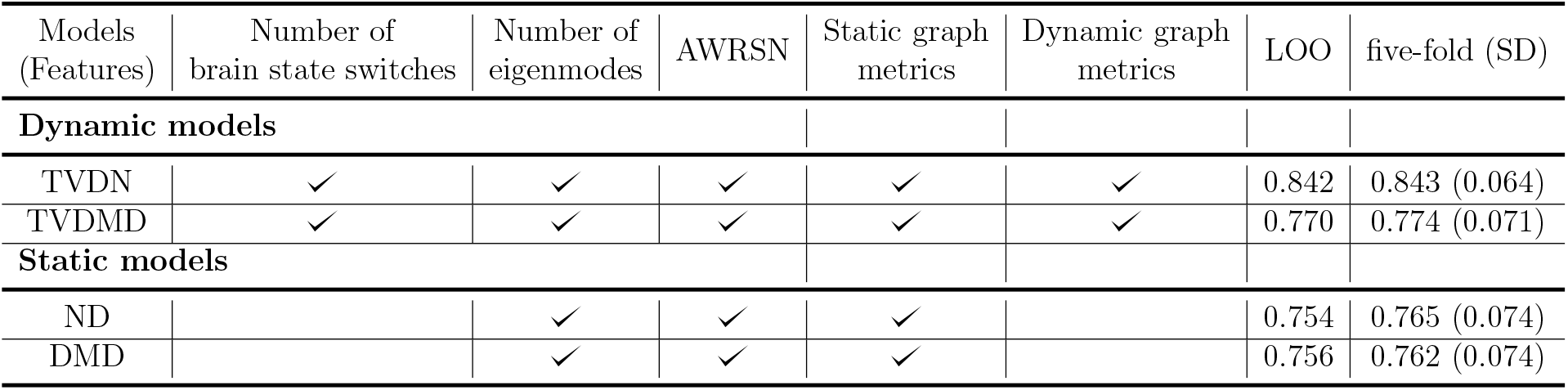
The features and graph metrics selected and the corresponding AUCs of TVDN, TVDMD, ND and DMD models. Static graph metrics include *L*mean, *Q*mean and the dynamic metrics include *L*var, *Q*var. TVDN model performs better than other models with 84% AUC. LOO: leave-one-out; five-fold: five-fold cross validation; SD: the standard deviation of the AUCs over 10000 cross validations.

Finally, we study the effects of features and graph metrics from the benchmark TVDMD, ND and DMD models in distinguishing AD and control samples through a ridge regularized logistic regression. The predictors of interest are summarized in Table 2 and the results are summarized in Table 3. As shown in Table 3, the numbers of eigenmodes from all models have a significant positive effect in distinguishing AD and control subjects. However, the effect size of the number of eigenmodes from the TVDMD model is much smaller than those from the TVDN, ND, DMD models. Furthermore, the effect size of the number of brain state switches from the TVDMD model is much smaller than those from the TVDN model. Moreover, consistent with the finding in Section 2.2, none of the graph metrics from the static models has significant effect on AD and control classification.

**Table 3:** TVDMD, ND, DMD results from a ridge regularized logistic regression: the effects from the predictors of interest on the AD and control classifications. The predictors include the number of eigenmodes, number of brain state switches, the dynamic and static graph metrics and the AWRSN over 68 regions of interest. Only the effects of the scalar predictors are presented. All predictors are all standardized by their sample means and standard deviations.

| Features and graph metrics | Effect | 95% CI | P-value |
| --- | --- | --- | --- |
| TVDMD scalar features and graph metrics |  |  |  |
| Number of eigenmodes | 0.008 | ( 0.002,0.014) | 0.005 |
| Number of brain state switches | 0.010 | ( 0.004,0.015) | 0.001 |
| $L_{\text{var}}$ | 0.010 | ( 0.004,0.015) | 0.001 |
| $Q_{\text{var}}$ | 0.008 | ( 0.003,0.014) | 0.004 |
| $L_{\text{mean}}$ | -0.005 | (-0.010,0.001) | 0.115 |
| $Q_{\text{mean}}$ | -0.000 | (-0.006,0.006) | 0.980 |
| ND scalar features |  |  |  |
| Number of eigenmodes | 0.104 | ( 0.042, 0.167) | 0.001 |
| $L_{\text{static}}$ | -0.011 | (-0.087, 0.065) | 0.775 |
| $Q_{\text{static}}$ | 0.040 | (-0.035, 0.115) | 0.293 |
| DMD scalar features |  |  |  |
| Number of eigenmodes | 0.131 | ( 0.072, 0.189) | < 0.001 |
| $Q_{\text{static}}$ | -0.063 | (-0.132, 0.006) | 0.074 |
| $L_{\text{static}}$ | -0.022 | (-0.097, 0.052) | 0.554 |

### 2.5 The relationship between TVDN features and the graph metrics

We further study the effects of TVDN scalar features on the brain network connection through examining their correlation with the graph metrics. This analysis aims to facilitate the interpretation of the TVDN features and reveal the graph information that explained by the TVDN features. We study the Pearson’s correlation between each of the two TVDN scale features and each of the eight graph metrics. Here all 16 variables are standardized through dividing by their sample standard deviations. We use p-value*<* 0.003 (0.05*/*16) as a significance cutoff after adjusting for multiple testing. It is expected that all the dynamic graph metrics in Table 4 are positively correlated with the frequency of brain state switching, because a larger number of brain state switches implies higher dynamicity in the FC which leads to more variable graph metrics. Furthermore, *Q*_mean_ and *Q*_max_ also are significantly correlated with the number of eigenmodes suggesting that a smaller number of eigenmodes implies denser connections within the subdivisions of the brain network from the dynamic FC. Moreover, the number of eigenmodes is negatively correlated with *L*var, suggesting that a greater number of eigenmodes implies less variations of path length over time. Finally, since the static model does not contain dynamic information, the TVDN scalar features are not associated with the graph metrics from the static model.

**Table 4:** Pearson’s correlation between TVDN features and graph metrics. *L*var, *Q*var are the variances of the standardized path length, and modality over time; *L*_mean_, *Q*_mean_ are the means of corresponding graph metrics over time; *L*_max_, *Q*_max_ are the graph metrics in the segments with the maximal dwell time; and *L*_static_, *Q*_static_ are the graph metrics from the static FC model. p-value*<* 0.003 is used as the significance cutoff to account for multiple testing.

| TVDN features | Number of eigenmodes |  | Number of brain state switches |  |
| --- | --- | --- | --- | --- |
| Graph metrics | Correlation | p-value | Correlation | p-value |
| TVDN dynamic graph metrics |  |  |  |  |
| $L_{\text{var}}$ | -0.268 | < **0.001** | 0.234 | **0.002** |
| $Q_{\text{var}}$ | -0.120 | 0.114 | 0.284 | < **0.001** |
| TVDN static graph metrics |  |  |  |  |
| $L_{\text{mean}}$ | -0.170 | 0.024 | -0.014 | 0.853 |
| $Q_{\text{mean}}$ | -0.371 | < **0.001** | 0.032 | 0.670 |
| $L_{\text{max}}$ | -0.111 | 0.143 | -0.018 | 0.811 |
| $Q_{\text{max}}$ | -0.254 | **0.001** | -0.034 | 0.656 |
| Graph metrics from the static model |  |  |  |  |
| $L_{\text{static}}$ | -0.160 | 0.034 | 0.225 | 0.003 |
| $Q_{\text{static}}$ | -0.200 | 0.008 | 0.011 | 0.880 |

## 3 Discussion

We demonstrate, for the first time, that the number of brain state switches, representing the temporal complexity of the brain network, in high-temporal resolution resting state MEG extracted by the time-varying dynamic network (TVDN) algorithm is an important feature that predicts AD. Specifically, AD subjects have fewer brain state transitions and in turn longer dwell periods in any given brain state than the control subjects. The number of eigenmodes, representing the spatial complexity of the brain network, is also an important predictor for AD, where AD subjects have greater number of eigenmodes, and hence a more heterogenous and complex functional brain network structure. We also demonstrate that the variability of graph metrics such as path length and modularity of dynamic functional connectivity periods are reduced in AD subjects. Interestingly, the static graph metrics corresponding to the brain state with the maximal dwell time or the mean of all brain states are not distinguishable between AD and control groups, while the dynamic graph metrics that correlated with the brain state switching are significantly different in the two groups. Using a data driven approach, TVDN identifies AD associated spatial features, that are different between AD and controls, in the brain regions that reflect high tau accumulation. When compared with predictions using features from other dynamic and static FC benchmarks, we show that features from TVDN leads to the best sensitivity and specificity for distinguishing AD and control samples. These results highlight the importance of dynamic functional connectivity in resting-state data for understanding the neural pathophysiology of AD.

### High resolution MEG data provide convincing evidence that the brain state switching patterns are altered in AD

Although static FC features extracted from MEG activity have proven to be reliable across different MEG laboratories (Geisseler et al., 2016) and have demonstrated to be an early biomarker of AD burden (Bajo et al., 2012; Fernández et al., 2006), literature is still scarce in studying the dynamic features from MEG data for neurodegeneration. Our study utilizes the novel TVDN method to extract the dynamic functional connectivity and brain state transitions from MEG data. We find that the minimal dwell time in the brain states is around 2 seconds, and half of the dwell times are less than 5 seconds. Such fast brain state transitions are difficult to capture by using low time resolution image modalities such as fMRI, because the samples within each stationary time segment are too limited to provide accurate estimates of the high dimensional whole brain functional connectivity. On the contrary, high resolution MEG data provide sufficient samples in each time segment to estimate functional connectivity, and hence can be used to capture fast brain state transitions that distinguish the AD and control samples.

### TVDN graph metrics are highly associated with AD and control classification

The application of graph theory to static resting state functional connectivity in AD has provided conflicting results. Based on the resting state fMRI data, Supekar et al. (2008) show no difference in the average path length between the AD and control samples in their study, while Sanz-Arigita et al. (2010) show a decreased average path length in the AD patients compared to the healthy control. In a MEG study, Stam et al. (2009) show that the similarities of the graph metrics between AD and control samples varies across different frequency bands. Our results show that the graph metrics from the static models cannot effectively distinguish AD and control samples, while Figure 3 shows that the variations of the graph metrics provide important information to distinguish AD and control subjects, which can only be obtained from dynamic FCs. Furthermore, the graph metrics in the control group generally have higher variability than those in the AD group and the variability patterns of the path length and modularity in AD and control groups are significantly different. These results highlight the importance of considering dynamic FC in AD studies.

### TVDN features sufficiently capture the information in the graph metrics

A comprehensive study on the TVDN features and graph metrics in Section 2.5 finds that the TVDN features are strongly associated with the graph metrics that differentiate AD and control groups (*L*var, *Q*var). For example, the number of eigenmodes negatively affects *L*var, suggesting that a greater number of eigenmodes implies less variations of path length over time. In addition, the number of brain state switches are positively associated with the dynamic graph metrics. Finally, when adjusting for the TVDN features, graph metrics do not contribute to the AD and control classification (Table 1), which indicates that the information in the graph metrics regarding the AD and control distinction is sufficiently captured by the TVDN features.

### AD patients have larger number of eigenmodes implying higher spatial complexity of the brain network

Figure 2 shows that the AD patients have a significantly larger number of eigenmodes, which implies a higher spatial complexity on average when comparing with the healthy controls. This is because the additional eigenmodes in the AD subjects introduce new bases in the lower dimensional manifold, which results in a more heterogenous and complex network structure. This is also supported by the fact that a higher number of eigenmodes is negatively associated with a lower modularity (Table 4), which leads to higher structural complexity (Baldwin and Clark, 2000; Sinha et al., 2018). These additional eigenmodes can form up AD specific pathological networks that do not exist in healthy subjects. Consistent with this idea, increase prevalence of pathophysiological epileptiform activity network structures have indeed been reported in AD when compared to healthy controls (Ranasinghe et al., 2022a; Vossel et al., 2016).

### Spatial patterns of dynamic connectivity changes overlap with the regional anatomy of AD pathophysiology

Figure 4 (a) shows the regional patterns of the estimated effects from AWRSN that distinguish AD from healthy elderly individuals. These regions include inferior and posterior temporal cortices and posterior parietal-occipital cortices, which reflects the same regional distribution of high tau accumulations, earliest hypometabolism and go onto develop greatest neuronal loss in patients with AD (Jagust, 2018). Distributions of tau accumulation both in space and time have been linked to network connectivity measures using various static network features, where functional connectivity based models could reliably predict individual variability of tau accumulation in AD (Franzmeier et al., 2020b,a). The proximity of spatial patterns between abnormal dynamic functional connectivity indices and AD pathophysiology relevant regional anatomy suggests that dynamic functional connectivity features, in addition to static features may also be worthwhile indices to explore as additional, complimentary predictors of AD pathophysiology. It is also noteworthy that the spatial distribution of dynamic connectivity differences is more left predominant in our findings. Although the biological significance of this lateralization of dynamic functional changes is yet to be explored, such asymmetry has been observed in previous resting state functional connectivity studies as well (Medvedev, 2014; Di et al., 2014). These results encourage further exploration of brain transitions within and between the two brain hemispheres.

### Dynamic features from TVDN with MEG imaging enhances the sensitivity and specificity of AD and control classifications

We compare prediction accuracies of using TVDN features, the features from TVDMD, a sliding window based method, and the features from static FC models. As shown in Table 2 and Figure 4 (b), TVDN has much higher prediction accuracy than the other models do, which shows the superiority of TVDN on extracting robust features that are highly differential in AD and control samples. Furthermore, the prediction accuracies are improved by introducing the temporal features from TVDN, the number of brain state switches, into the prediction model. This can be seen from Table 2 and Figure 4 (b) that using the TVDN features yields the highest sensitivity and specificity in classifying AD versus control subjects. Moreover, the results in Table 1 and Table 3 show that the effect size of the number of eigenmodes and the number of brain state switches from TVDN are much larger than those from the TVDMD model. Furthermore, using the features from the TVDN model gives the best prediction accuracy among all the comparative methods. The lower prediction accuracy and smaller effect sizes from the TVDMD method could attribute to its deficiencies inherited from the sliding window based method, such as the arbitrariness of the window length selection and the curse of dimensionality. Collectively, these results indicate that TVDN is a more reliable method to detect brain state switches than TVDMD.

### fMRI studies of dynamic FC in AD

The present study uses MEG imaging to demonstrate that dynamic functional connectivity features are abnormal and have predictive value in AD. Here, we review a larger fMRI literature on dynamic functional studies in AD. Consistent with our dwell time findings, Jones et al. (2012) suggest that the dwell time in the default mode network are distinctive between AD dementia patients and healthy controls. Brenner et al. (2018) also show that amnestic MCI patients spend more time in a single dominant state. Also consistent with our observation of increased number of switches in AD, Córdova-Palomera et al. (2017) show decreased global metastability between functional states when comparing the patients with mild cognitive impairment (MCI) and healthy controls. Similar to our findings on the predictablity of dynamic spatial features, Fu et al. (2019) examine the shared and specific dynamic functional connectivity in subcortical ischemic vascular disease and AD. Dautricourt et al. (2022) show that dynamic FC states are differentially associated with dementia risk. However, the above mentioned studies have not clearly demonstrated that dynamic FC features can distinguish AD patients from healthy controls (Jones et al., 2012; Córdova-Palomera et al., 2017; Brenner et al., 2018). In contrast, a recent study by de Vos et al. (2018) reveals higher accuracy to distinguish AD dementia from healthy controls using the variability of FC across time as a feature than static FC features, perhaps the first clear evidence that the dynamic FC can be a strong predictor of AD. However, their prediction model included a large number of features and did not fully address which dynamic FC features were important to distinguish AD and healthy subjects (de Vos et al., 2018). Therefore, whether dynamic FC features have predictive power to distinguish AD patients, and if so which features are important to drive these predictions remain unknown from these prior fMRI studies. Extending TVDN to fMRI data is important to address these questions. Collecting resting state fMRI data and further research along these lines are ongoing in our laboratory.

### Dynamic FC features have predictive value in AD

It must be borne in mind that the present classifier results by themselves do not argue for the exclusive use of dynamic FC features as predictors of AD, whether for diagnostic or prognostic purposes. Indeed, under the AUC metric of classifier performance, we have reported close to 84% accuracy, which is good but not accurate enough for diagnostic purposes. Therefore, the current results must be interpreted as supportive of a valuable role for dynamic FC features in a diagnostic application, perhaps in combination with static FC features, as well as other imaging biomarkers like regional atrophy and molecular PET imaging. Such applications will not only be able to correctly predict AD diagnosis, but the addition of dynamic FC will have imbued the application with a hitherto unfeasible ability to differentiate between various AD subtypes. Our contribution is to show that a model-based TVDN approach provides far more predictive power in the use of dynamic FC compared to alternate means of obtaining dynamic FC features, and that TVDN-derived dynamic features uncover important processes of the AD pathophysiology that are currently being unreported by conventional static FC methods.

Arguably, a far more clinically important use case for proposed approach will be as a means to understand how AD affects the dynamicity of the brain, and in future work, to explain how these dynamic changes may be caused by underlying pathology (tau, amyloid and hypometabolism) and the resulting network disconnection. For instance, what specifically about the deposition of tau and amyloid in and around neurons could cause an increase in dwell times in the AD brain, indicative of its brain states being “stickier” than normal brains with no pathology? Although out of scope of current work, such studies would require new models that relate pathology to functional dynamics, and would provide valuable new insights into AD pathophysiology that is currently lacking.

## 4 Methods

### 4.1 Data and preprocessing

Each participant in our study underwent a complete clinical history, physical examination, neuropsychological evaluation, brain magnetic resonance imaging (MRI), and a 10-min session of resting MEG. All the acquisition and processing pipelines are the same as that for a previous study (Ranasinghe et al., 2022b). All participants were recruited from research cohorts at the University of California San Francisco-Alzheimer’s Disease Research Center(UCSF-ADRC). Informed consent was obtained from all participants and the study was approved by the Institutional Review Board (IRB) at UCSF (UCSF-IRB 10-02245). Demographics of cohorts are summarized in Table 5.

**Table 5:**
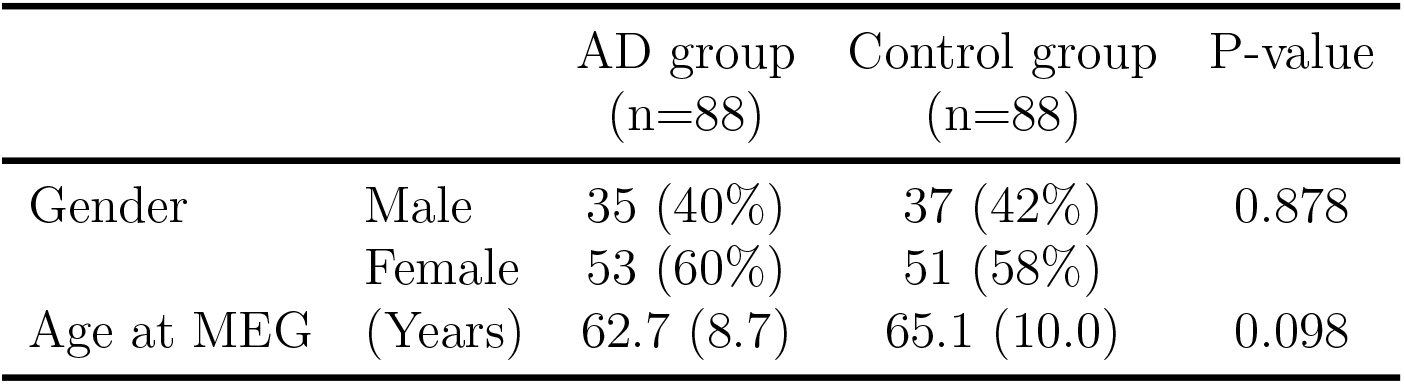
Baseline characteristics: The number of samples (percentiles) for gender and the mean (standard deviation) for age are presented

### 4.2 Resting state MEG data acquisition and preprocessing

MEG scans were acquired on a whole-head biomagnetometer system (275 axial gradiometers; MISL, Coquitlam, British Columbia, Canada), following the same protocols described previously (Ranasinghe et al., 2020, 2022b). Each subject underwent MEG recordings at rest, eys-closed and supine for 5–10 min. Three fiducial coils including nasion, left and right preauricular points were placed to localize the position of head relative to sensor array, and later coregistered to each individual’s respective MRI to generate an individualized head shape. Data collection was optimized to minimize within-session head movements and to keep it below 0.5 cm. 5–10 min of continuous recording was collected from each subject while lying supine and awake with eyes closed (sampling rate: 600 Hz). We selected a 60s (1 min) continuous segment with minimal artifacts (minimal excessive scatter at signal amplitude *<*10 pT), for each subject, for analysis. The study protocol required the participant to be interactive with the investigator and be awake at the beginning of the data collection. Spectral analysis of each MEG recording and whenever available simultaneously collected scalp EEG recordings were examined to confirm that the 60-second data epoch represented awake, eyes closed resting state for each participant. Artifact detection was confirmed by visual inspection of sensor data and channels with excessive noise within individual subjects were removed prior to analysis.

### 4.3 Source space reconstruction of MEG data and spectral power estimation

Tomographic reconstructions of the MEG data were generated using a head model based on each participant’s structural MRI. Spatiotemporal estimates of neural sources were generated using a time–frequency optimized adaptive spatial filtering technique implemented in the Neurodynamic Utility Toolbox for MEG (NUTMEG; https://nutmeg.berkeley.edu/).

To prepare for source localization, all MEG sensor locations were coregistered to each subject’s anatomical MRI scans. The lead field (forward model) for each subject was calculated in NUTMEG using a multiple local-spheres head model (three-orientation lead field) and an 8-mm voxel grid which generated more than 5000 dipole sources, all sources were normalized to have a norm of 1 (Dalal et al., 2008, 2011). The source space reconstruction approach provided amplitude estimations at each voxel derived through the linear combination of spatial weighting matrix with the sensor data matrix (Dalal et al., 2008). A high-resolution anatomical MRI was obtained for each subject (see below) and was spatially normalized to the Montreal Neurological Institute (MNI) template brain using the SPM software (http://www.fil.ion.ucl.ac.uk/spm), with the resulting parameters being applied to each individual subject’s source space reconstruction within the NUTMEG pipeline (Dalal et al., 2011). We then derived regional power spectra based on Desikan–Killiany atlas parcellations for the 68 cortical regions depicting neocortex and allocortex, the latter including the entorhinal cortex. Regional power spectra were obtained by averaging the spectra of vowels within each region and then converted to dB scale.

### 4.4 Time-varying dynamic network model

The time-varying dynamic network (TVDN) model is a robust method to extract dynamic FCs from the neuroimaging data (Jiang et al., 2022), which assumes the brain states experience discrete and discontinuous changes over time. The dynamic FC features contain static spatial feature (RSNs) and dynamic temporal feature (the dynamic weights of the RSNs).

Let **X**(*t*) be a *d* dimensional vector, denoting the brain activity at time *t* at *d* numbers of ROIs, and let **X**^*′*^(*t*) be its derivative. The TVDN model assumes

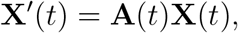

where **A**(*t*) is a *d* × *d* connectivity matrix. The connectivity matrix **A**(*t*) can be further decomposed as **A**(*t*) = **UΛ**(*t*)**U**^*−*1^, where the columns of **U** represent the static RSNs, and the eigen-value matrix **Λ**(*t*), varying across the time, represents the importance of each RSN. The real and imaginary parts of **Λ**(*t*) represent the growth constant and signal frequency, respectively. Since only a small number of RSNs are operational in the brain, typically ranges from 7-20 (Yeo et al., 2011), **A**(*t*) is assumed to be a low-rank matrix, where the rank of the **A**(*t*) represents the number of distinct static RSNs that are active in the resting state.

To extract the dynamic FC features, we first perform a B-spline smoothing step to obtain noise free versions of **X**^*′*^(*t*) and **X**(*t*). Then, we implement a kernel regression step to obtain the Nadaraya–Watson estimator (Nadaraya, 1964; Watson, 1964) of **A**(*t*), denoted by 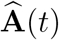. The static spatial feature **U** is then extracted as the top *r* eigen-vectors of 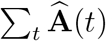, where we choose rank *r* so that the magnitude of the top *r* eigen-values of 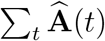 comprises 80% of the total sum of the magnitude of them. Next, the brain state switches are detected by minimizing a modified Bayesian information criteria (MBIC) through the dynamic programming algorithm (Jackson et al., 2005) based on a low dimensional trans-formation 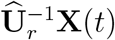, where 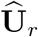 is the first *r* columns of the estimated **U**, and 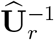 is the *r* × *d* dimensional generalized inverse of 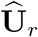. The brain activities are separated to different segments after the detection procedure. Then we refit TVDN model in each segment, while assuming **A**(*t*) is a constant. Furthermore, the temporal dynamic weighted are obtained as the eigen-values of estimated **A**(*t*) in each stationary segment, denoted by 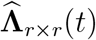, which is a *r* × *r* dimensional matrix. We obtain the WRSN feature as the column sum of 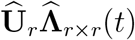. Finally, we first obtain 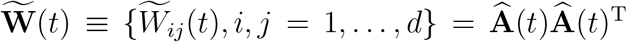 and construct the functional connectivity matrix through the following two steps:

1. let 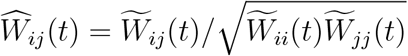
2. perform fisher transform and obtain the functional connectivity matrix **W**(*t*) with the *i, j*th entry to be 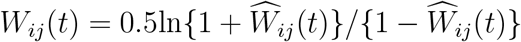

All the tuning parameters in the TVDN procedure are tuned based on the control samples.

The TVDN analysis is performed using the Python code from https://github.com/feigroup/TVDN, which contains detailed documentation of the code usage.

### 4.5 Dynamic mode decomposition model and network diffusion model

We implement the dynamic mode decomposition model (DMD) and the network diffusion model (ND) for comparison.

The DMD model assumes **X**(*t* + 1) = **AX**(*t*) (Brunton et al., 2016), where **A** is a *d* × *d* constant matrix. We obtain the estimated **A** by minimizing the sum of the squared distance between **X**(*t* + 1) and **AX**(*t*) over time. Then we obtained the estimated RSNs, denoted by 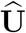, as the first *r* eigen-vectors corresponding to the top eigen-values, whose summation comprises 80% of the total sum of the eigen-values. Furthermore, we construct the WRSN feature as the column sum of 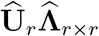 where 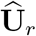 is a *d* × *r* dimensional eigen-vector matrix, and 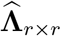 is a *r* × *r* dimensional eigen-value matrix. Since there is only one segment resulted from the DMD model, AWRSN feature is the same as the WRSN feature. Finally, we obtained the 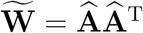 and use steps 1, 2 in Section 4.4 to generate the final connectivity matrix **W**.

The ND model assumes **X**^*′*^(*t*) = **AX**(*t*). We obtain the estimated **A** by minimizing the sum of the squared distance between **X**^*′*^(*t* + 1) and **AX**(*t*) over time. The remaining steps for extracting spatial features from the ND model are the same to the DMD model.

We implement TVDMD as follows where we construct windows with 192 frames sliding by 24 frames in each step. In each sliding-window, TVDMD extracts the dynamic modes (Brunton et al., 2016; Kunert-Graf et al., 2019) from the brain signals, and obtains the WRSN in each sliding windows using the same procedures as those described in DMD model above. The selection of the window size and step size leads to 292 number of sliding windows, which is similar as those used in Kunert-Graf et al. (2019). We then use Kmeans algorithm to cluster the WRSNs to 3 clusters, which is the average number of switches from the control group with the TVDN method. We define a switch point as the time where there is a cluster membership change before and after the time. These switch points divide the MEG data into multiple segments, and in each segment we reestimate the WRSN and the number of eigenmodes, whose corresponding magnitude of the eigenvalues comprises 80% of the total sum of the magnitude of the eigenvalues. We then average the WRSN as the AWRSN and calculate the maximal number of RSNs across the multiple segments as the number of RSNs.

The DMD and ND analysis were conducted using the Python code (https://github.com/feigroup/TVDN-AD).

### 4.6 Graph metrics

The brain networks extracted from functional connectivity models can be represented by graphs, which are the combination of ROIs, the nodes in the graph, and the edges (Boccaletti et al., 2006), the region-wise connections in the graph. The strength of the connections among the ROIs, namely the edge weights, are mathematically captured by the entries of the functional connectivity matrix **W**.

The path length *L* of the graph is defined as

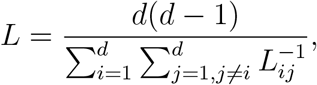

where *L*_*ij*_ is the shortest length of the path that goes from regions *i* to *j*, where the length of a path is the summation of the inverse of edge weights on the path (Wang et al., 2013).

The modularity *Q*_max_ is a statistic that quantifies the degree to which the network can be subdivided into different groups (Newman, 2006), where the optimal group structure is obtained by maximizing the number of within-group connections, and minimizing the number of between-group connections.

Let *W*_*ij*_ be the *i, j*th entry of the functional connectivity matrix, representing the strength of connection between ROI *i* and ROI *j*. If *W*_*ij*_ ≠ 0 (*W*_*ij*_ = 0), region *i* and *j* are connected (disconnected). For a given partition *p* of the graph, the modularity index 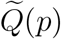 is defined as

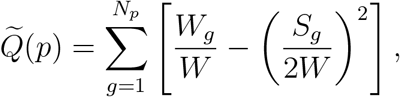

where *N*_*p*_ is the number of groups in the partition, *W* = Σ_*i,j*_ *W*_*ij*_ and *W*_*g*_ is sum of all the edge weights between the regions in the group *g*. Here let 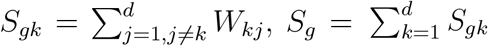 is the sum of the nodal strength in group *g*. Finally 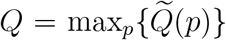, which is obtained by using the spectral algorithm described in (Newman, 2006).

To eliminate this dependence of the graphic features on the size of the graph, we normalize the three features through dividing them by an ensemble of 100 random networks, which is described as follows. We randomly permute the values in **W**, and create 100 sets of pseudo graph features. We them divide the each of the two graphic features by the corresponding means of the pseudo graph features over the 100 permutations.

To extract the graph characteristics, we adopted the bctpy package in python (https://pypi.org/project/bctpy/), which contains the detailed documentation of the code usage.

### 4.7 Logistic regression with ridge regularization

In our manuscript, we adopted the logistic regression with the ridge penalty as the classifier for the AD and control groups. The ridge regularization was utilized due to the high-dimension feature in our regression (Hoerl and Kennard, 1970).

With the ridge penalty, the loss function to optimize becomes

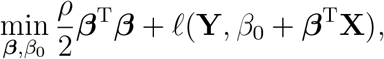

where **X** are the FC features, which can be different from TVDN, DMD, ND models, **Y** = (*Y*_1_, …, *Y*_*N*_)^T^ is a vector a binary indicator with *Y*_*i*_ = 1 or *Y*_*i*_ = 0 if the *i*th subject is a AD patient or a healthy control, respectively. The function *ℓ*(*·*) is the negative log likelihood of the logistic regression and *ρ >* 0 is the penalty parameter. The penalty parameter is tuned by the leave-one-out evaluation within the training set to ensure no testing data information is used during the training procedure.

For each model, we implemented the two ridge regularized logistic regressions. We use first ridge regression to select significant predictors as the ones whose estimated 95% confidence intervals do not cover zero. We then utilized the selected important predictors in the second ridge regression model to evaluate the sensitivity and specificity of classifying AD and control samples through leave-one-out (LOO) and five-fold cross prediction over 10000 repetitions. The penalty parameters are tuned based on the training data to avoid information leaking in the prediction.

We implemented the logistic regression with the ridge penalty by the LogisticRegression function in sklearn package in Python. The detailed documentation of the package is accessible at https://scikit-learn.org/stable/modules/generated/sklearn.linear_model.LogisticRegression.html.

